# Invasive plants as novel food resources, the pollinators’ perspective

**DOI:** 10.1101/018952

**Authors:** Ignasi Bartomeus, Jochen Fründ, Neal M. Williams

**Affiliations:** Estación Biólogica de Doñana - Consejo Superior de Investigaciones Científicas (EBD-CSIC), Dept. Integrative Ecology. Sevilla, Spain. < >; University of Guelph, Dept. Integrative Biology, Guelph, ON, Canada; University of California, Dept. Entomology and Nematology, Davis, CA, USA

## Abstract

Biological invasions are one of the main drivers of global change and have negatively impacted all biomes and trophic levels (Hobbs 2000, Vilà et al. 2011). While most introduced species fail to establish, or establish small naturalized populations (hereafter exotic species), a few become invasive and rapidly increase in abundance and/or range (hereafter invasives or invaders; Pysek et al. 2004). It is these invader species that are most often linked to negative impacts on native / endemic communities. Although most interactions between invasive and native species at the same trophic level result in negative direct impacts (e.g. plant-plant competitive interactions, Vilà et al. 2011), when the invasive plant species can be used as a resource those interactions can also be positive for consumers such as native herbivores, predators or mutualists, at least for some species (Heleno et al. 2009, Bezemer et al. 2004). Entomophilous exotic plant species, for example, not only compete directly for space and light with other plants, but also offer resource opportunities for the native pollinator community (Stouffer et al. 2014). Most research on this topic to date has taken the plant perspective, focusing on how successful plant invaders integrate into the native plant-pollinator interaction networks (Vilà et al. 2009), and how this integration in turn impacts the native plant community (Morales and Traveset 2009). However, species specific responses of pollinators to the addition of exotic plants are rarely taken into account. This represent an important gap in our knowledge as pollinator foraging choices determine the structure of interactions within communities, which in turn have important implications for the community stability (Thébault and Fontain 2010) and functioning (Thomson et al. 2012). How different pollinators respond to the changed composition of floral species within the community that results from exotic plant invaders deserves more attention.

---

From the pollinators’ point of view, exotic plants are novel food resources, and as such their relative abundance, attractiveness, rewards (i.e. nutritional value) and distinctiveness partly determine their use by various pollinators (Carvalheiro et al. 2014). Although exotics as a group are not preferred or avoided within their new communities (Williams et al. 2010), it might be that particular exotic species are preferred by some pollinators while avoided by others. The intrinsic preferences for different plant hosts is an important factor determining host use. Hence, the direct benefits or costs of a novel resource use will differ among pollinator species. Moreover, in a community context, the preferences of each pollinator affect the other pollinators’ choices, potentially leading to indirect effects. For example, some pollinator species may obtain indirect benefits if the invasive plants distract other pollinators from natives, reducing competition. Alternatively, pollinators may pay indirect costs if competition is increased or if invasive plants reduce the availability of a preferred native plant.

We review the evidence on direct and indirect benefits and costs of invasive plants on pollinators and re-analyze Williams et al. (2010) dataset on pollinator specific preferences so as to identify species that prefer some exotic plants over native plants and vice-versa. This information is crucial to understanding the consequence for the pollinator community because if only some pollinators take advantage of alien plants this can favour populations of some pollinator species (winners) over others (losers). By using an approach that takes into account both pollinator behavioral responses and interaction network structure, we can better understand the invasion process, with important implications for conservation actions.

## Effects of plant invasions on pollinator populations and community structure

The impact of invasive plants interacting with native pollinators has received considerable attention for its potential to disrupt native mutualisms (Traveset and Richardson 2006). However, most work to date has focused solely on how invasive plant interactions with native pollinators changes the pollination success of native plants. Interestingly, existing data show that invasive plants can have positive, neutral, or negative effects on pollination of native plants (Bjerknes et al. 2007). The contrasting results may reflect invasive plant density (Muñoz and Cavieres 2008, Dietzsch et al. 2011), spatial aggregation (Cariveau and Norton 2009) or flower morphology and attractiveness (Morales and Traveset 2009, Carvalheiro et al. 2014).

In contrast, the effects of exotic or invasive species on the pollinator populations and communities have received far less attention (Stout and Morales 2008). Because most plant-pollinator systems are generalized (Waser et al. 1996), invasive plants are usually well integrated in the plant–pollinator network of interactions (Vilà et al 2009). Hence, we might expect overall effects on pollinators to mirror the changes (positive or negative) in floral resources offered by the newly invaded community. If the entomophilous exotic or invasive plants increase the resources present in the community, this should also allow the increase of most pollinator populations (Tepedino et al. 2008). Stout and Morales (2008) cite indirect evidence that some social pollinators (e.g., bumblebees) can be favoured by non-native mass flowering crops (Westphal et al. 2003, Herrmann et al. 2007), which may be analogous to the effect of abundant invasive species. However, the same authors note examples where exotic plants are not used by native pollinators due to flower morphology or chemistry. Despite Stout and Morales’ (2008) call for more research on this topic, few additional studies have been published since then.

Current evidence suggests that food resource availability may indeed regulate pollinator’s populations, at least those of bees (reviewed in, Roulston and Goodell 2011, Williams and Kremen 2007, Crone 2013, but see Steffan-Dewenter and Schiele 2008 for potential regulation by nesting resources). However, studies of the effect of exotic plants on pollinators’ population dynamics are extremely rare, particularly for non-invasive exotic plants that establish small populations. Palladini and Maron (2014) provide one of the few examples of effects of exotic plants (mainly *Euphorbia esula*) on the reproduction of a solitary bee species (*Osmia lignaria)*. For this species, the number of nests established and offspring production per female was positively related to native plant abundance and negatively related to exotic plant species. This may be because although *Osmia lignaria* foraged on exotics for nectar, the species did not use *Euphorbia esula* exotic pollen to provision nests. Thus, the specific ability to use the invader resources emerges as a key factor affecting the potential impacts of the invader on pollinators. The only other evidence to date for direct effects of invasive plants through resource augmentation is for bumblebees, whose annual social life history allows demographic responses to be measured within a single season. Within-season abundance can increase almost four times in sites invaded by the plant *Lupinus polyphyllus*, compared to in non-invaded sites (Jakobsson and Padrón 2014). In addition, the foraging season of *Bombus terrestris* in the United Kingdom can be extended into winter through its use of exotic plants that fill a late season phenological niche (Stelzer et al. 2010). Such within-season demographic responses are likely to have longer-term population effects, although no study has quantified such effect to date.

A second group of studies provides indirect evidence that exotic plants affect pollinators’ by using community approaches to compare invaded with non-invaded sites. These studies show a variety of pollinator responses, including increased abundance and species richness in invaded sites (Lopezaraiza-mikel et al. 2007), lower abundance and diversity of pollinators in invaded areas (Moron et al. 2009), or no difference in abundance between invaded and non-invaded sites (Nienhuis et al. 2009). It is therefore not surprising that a recent meta-analysis reported no changes in overall pollinator abundance in invaded sites (Montero-castaño and Vilà 2012). However, the studies included in the meta-analysis were not designed to infer population changes, and the result should be interpreted with caution. Moreover, most of the examples concern abundant invasive plants. Plant abundance can strongly influence pollinators decision to incorporate a new plant into its diet (Valdovinos et al. 2010), and thus the results may differ when examining non-invasive exotic plants. Likewise, given the wide array of pollinators ranging from birds and bats to bees and hoverflies, it is unlikely to find a consistent overall response.

## The importance of behaviour

While exotic plants can represent new resource opportunities for native animals, evidence suggests that only a minority of pollinator species can take advantage of these new opportunities. For example, generalist bees more commonly forage on invasive exotic plants than specialists (Lopezaraiza-Mikel et al., 2007; Tepedino et al., 2008; Padrón et al. 2009), or that social bumblebees are more common in invaded sites than solitary bees (Nienhuis et al. 2009). Bartomeus et al. (2009), for example, report that bees were more often recorded on native species than on the invader *Carpobrotus* aff. *acinaciformis*, except for the social bumblebee *Bombus terrestris* and for most beetles, which visited *Carpobrotus* flowers almost exclusively. Hence, pollinators can discriminate between native and exotic plants, and the decision of exploiting one or another can vary across species. Analyzing the invasion process as a novel resource availability for pollinators may give us a framework to predict which pollinators can beneficiate from the invasion process.

Incorporating any novel resource into the diet requires a series of conditions to be met. First, the pollinator must recognize the novel resource as a host, second, the visitor must be able to use this new resource and third, the resource must be profitable (i.e., a net benefit) for the visitor to exploit it. Hence, exotic plant use depends on intrinsic traits of the plant and pollinator. We cannot assume that for native pollinators exotic flowering species are always fundamentally different than the native flowering species; nevertheless, plants presenting new colors, shapes or chemical compounds may not be easily used by all pollinators in the community (Stout and Morales 2008). Furthermore many pollinators have innate preferences for certain colors and/or shapes (Gumbert 2000, Riffell et al. 2008), which may make novel flowers less attractive than the natives which have coevolved in the community. Neophobic responses also may make some pollinators unlikely to approach and explore novel food opportunities (Forrest and Thomson 2009). However, some pollinators may learn how to exploit new resources if their behaviour is flexible enough (Chittka et al. 2009, Forrest and Thomson 2009). In particular, bees, which are the main pollinator group both in numbers and in effectiveness (reviewed in Winfree et al. 2011), have a powerful neuronal system able to learn new tasks (Chittka et al. 2009) and their behavior flexibility has been suggested to be linked to their ability to persist in disturbed environments (Kevan and Menzel 2012). However the role of learning abilities in incorporating new foraging plants is little explored (Dukas and Real 1993) and most information comes from a handful of species (mostly social).

Even if the exotic plant is recognized as a potential host, pollinators might not be able to exploit the new plant species if they are unable to handle its flowers (Parker 1997, Corbet et al. 2001), or to digest its nectar (Adler and Irwin, 2005) or pollen (Sedivy et al. 2011, Palladini and Maron 2014). Morphological matching between flowers and pollinators may thus be important in determining pollinator visitation patterns (Gibson et al. 2012, Bartomeus 2013, Stang et al. 2009, see also Pearse et al. 2014 for antagonist insect-plant interactions). Second, even in the cases where pollinators recognize and can use the novel resource, their decision to include it in its diet will depend on its quality and abundance relative to others in the community. The thresholds for switching to a resource based on its quality or abundance have been show to be variable among different species in birds (Carnicer et al. 2008). Insect pollinators switch between foraging plants depending on the resource availability (Inouye 1978, Chittka et al 1997). Like for birds, thresholds for switching are likely to be different for different species, however, the experimental evidence that insect pollinators can discriminate between different resources and learn to forage on the preferred one is limited to a handful of bee species, and the switching strategies among species is mostly unknown.

## The importance of the community context

Pollinator-plant interactions do not occur in isolation, but form part of a complex network of interactions (Bascompte and Jordano 2007). For example, competitive exclusion between pollinator species can drive foraging behavior patterns (Jhonson and Hubell 1995). Invasive plant species can modify the native network structure not only by creating new interactions with native pollinators but also by modifying the existing plant-pollinator interactions among the native species (Bartomeus et al. 2008, Albrecht et al. 2014). Such changes are especially likely when invaders are abundant and provide accessible resources (i.e. nectar and pollen), thus potentially interacting with a large proportion of the resident pollinator species (Albrecht et al. 2014). In any case, modifications to pairwise interactions can have cascading effects throughout the community. For example, dynamic pollination network models have been used to show that removal of well-established invasive plants negatively affected the persistence of pollinator interactions through the network (Valdovinos et al. 2009, Carvalheiro et al. 2008). Detailed empirical studies, also show that co-flowering neighboring invasive species affect pollinator choices (Cariveau and Norton 2009, Waters et al. 2014).

The behavioural switching (also called interaction rewiring) between resources has been recently studied at the community level using network theory. There is increased evidence that pollinators can rewire their community-wide interactions depending on the context (Kaiser-Bunbury et al. 2010), such as the addition of a plant invasion. Jackobson and Padrón (2014) speculate that the attraction of bumblebees to the invasive plant *Lupinus polyphyllus*, reduced competition for the native plants, allowing an increase of solitary bee visits to natives. Similarly, Montero-Castaño (2014) showed that monopolization of the invasive species *Hedysarum coronarium* by honeybees allowed other bee species to establish interactions with natives that are not realized when the honeybee is present. Context dependent rewiring is supported by findings that despite for consistent species-specific preferences for certain flowers across communities, there is also important variation and flexibility in preferences among different contexts (Fründ et al. 2010). Hence, both direct effects and indirect effects on pollinators are expected after a plant invasion, and those can only be understood in a community-wide framework.

In Figure 1, we illustrate a simplified plant-pollinator network with two distinctive scenarios. In the first one (Fig. 1.B), we add an entomophilous exotic plant that does not reduce the abundances of other plants. Pollinator species able to use this plant (identified as p1 and p2) will establish new links with this exotic plant (blue links). Species p1 will have more food resources and can potentially increase its population over time (bigger blue circle denotes population size increase) whereas species p2 will experience a neutral effect because it changes from foraging on natives to foraging on the exotic. These are the direct (often neutral or positive) effects of the exotic plant on pollinators. Other pollinators may not be able or may not choose to visit the new invasive plant (p3), but as the competition for their preferred resources is changed, they may receive indirect benefits (p3). Experiments removing dominant pollinators have shown that a relaxed competition for resources may lead to diet expansion of some species (Brosi et al 2013), supporting our example with species p3.

**Figure 1.**
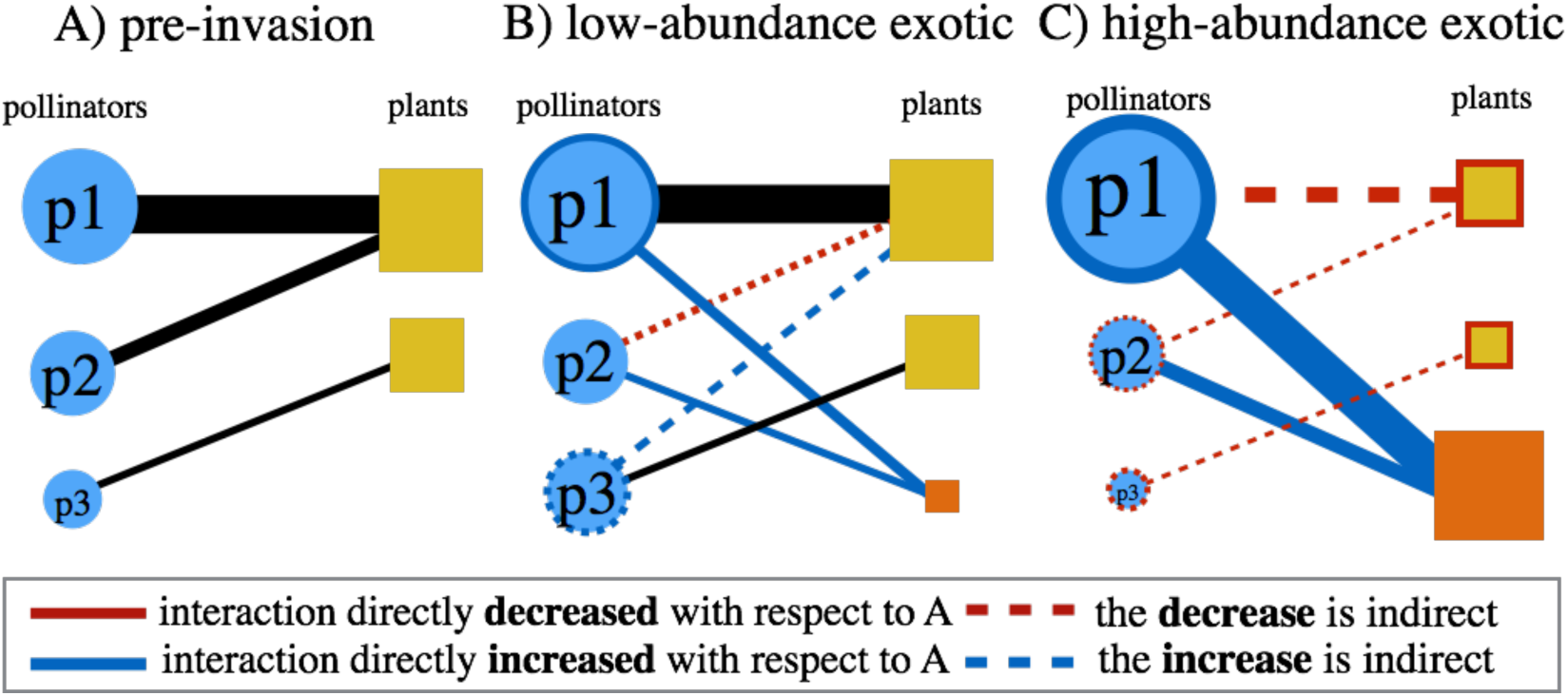
Simplified plant-pollinator networks before the invasion process (A) in two distinctive scenarios: (B) The exotic plant (orange square) adds a novel resource without affecting the rest of the community, and (C) a superabundant invasive plant (orange square) adds an abundant novel resource, while reducing native plants resource availability. In the schemes we can see the interactions established in each case (lines connecting pollinators and plants). The color of links depicts whether their frequency is increased (blue) or decreased (red) with respect to A. The changes in size of the plants or pollinators depict the winners (blue border) and losers (red border) of the invasion process. When the effect is indirect is noted with dashed lines. See text for further explanation.

In the second scenario (Fig 1. C), the exotic plant is an abundant invader that also reduces native plant abundance by direct competition (e.g. for space). In this scenario, only a few pollinators (p1) may benefit, while all others will experience increased competition for resources (p2, p3). This is an oversimplification, and of course the net benefit for pollinators will depend not only on the number of visits, but on the quality of those visits (e.g. reward uptake, nutritional content of the exotic species, etc.). Moreover, some species will require a variety of pollen sources to complete larvae development (Roulston and Cane 2000) highlighting the importance of maintaining plant diversity. The magnitude of the indirect and direct effects will depend also on the time-scale at which it is evaluated, with functional responses and local switching occurring faster than numerical responses (i.e. population growth). Moreover, the relative phenological timing of plants and bees can modify their mutual influence. All in all, the net costs and benefits are likely to depend on many factors, but this framework supports the scarce information presented above, where some social generalized species tend to increase their abundance after invasion by highly attractive species, but other pollinators have mixed responses.

## Case study: bee preferences in California

Can we predict which pollinators will be winners or losers of the invasion process? Measuring population responses or fecundity is a daunting task, especially at the community level; however, we can gain indirect evidence by looking at pollinator preferences. Within a plant community, pollinators do not prefer exotic plants as a group (Williams et al. 2010) or even prefer natives (Chrobock et al. 2013), but individual pollinator preferences have not been explored yet. Preference is defined as using a resource more than expected given its abundance. Conversely, avoidance occurs when a resource is underused relative to its abundance. The null model of no preferences is the case when pollinators visit flowers in proportion to their abundance in the community. Deviations from this null model can help us identify pollinator species that prefer exotic species (hence receiving a potential direct benefit) and species that avoid the exotics (hence, receiving negative, neutral or positive indirect effects in some cases). We recognize that we cannot infer direct fitness consequences, or predict indirect effects from a static network. Ours, nonetheless, is the first attempt to identify direct effects and serve as a proxy for identifying pollinator winners after the invasion process. Most importantly, this way we can emphasize that pollinators differ in their behavior, acknowledging that the effects on specific pollinators cannot be generalized. Furthermore, in the future, we can explore what determines pollinator preferences. Are they driven by plant traits, such as abundance or morphology, by pollinator traits, or a combination of both?

To explore this preference-based proxy, we used the same dataset used in Williams et al. (2010). For simplicity we show here only 7 sites from semi-natural habitats in California. This system is especially suitable to test our questions, because it contains several exotic plants, ranging from abundant invaders to naturalized exotic plants, as well as a variety of pollinator species. We calculated preferences pooling all sites, but we separate our analysis in three sampling periods (early, mid and late season). We treated periods separately because plant turnover was substantial over the season and otherwise might have masked the preference relationships.

First we re-evaluated that pollinators do not prefer exotic plants as a group within a quantitative framework, where expected (E) visitation values are calculated based on plant mean abundance across sites, and observed (O) visitation values are the sums of pollinator visitation to each plant across sites. Chi statistics were used for each of the three tests (i.e. one per season) to assess if there is an overall preference. The Pearson residuals of the Chi tests (O-E/√ E) estimate the magnitude of preference or avoidance for a given plant based on deviation from expected values and its significance was assessed by building Bonferroni confidence intervals (see Neu et al. 1974 and Byers and Steinhorst 1984). In order to test for differences between exotics and natives we compared the *Electivity* values of exotic and native plants using linear models. *Electivity* values (E’-O’/E’+O’, where E’ and O’ are the proportional expected and observed values, Ivlev 1964) are bound between −1 and +1, easier to interpret and highly correlated with Pearson residuals. R code to calculate these indexes can be found at (https://gist.github.com/ibartomeus/cdddca21d5dbff26a25e).

We show that when pooling visits for all pollinator species, some plants are preferred over others (Chi square test p-values for early, mid and late seasons < 0.001; *Electivity* values range from −0.84 to 0.82 indicating we find both over-preferred plants and under-preferred plants; Fig. 2). In accordance with the results reported in Williams et al. (2010), pooled pollinators showed no preferences for exotic plants in any season (Fig. 2; early season: F_1,25_ = 1.29, p-value: 0.27; mid season: F_1,22_ = 0.06, p-value: 0.81; late season: F_1,15_ = 0.46, p-value: 0.51), but this general trend does not contradict the fact that specific native or exotic plants are indeed preferred (in red in Fig. 2).

**Figure 2:**
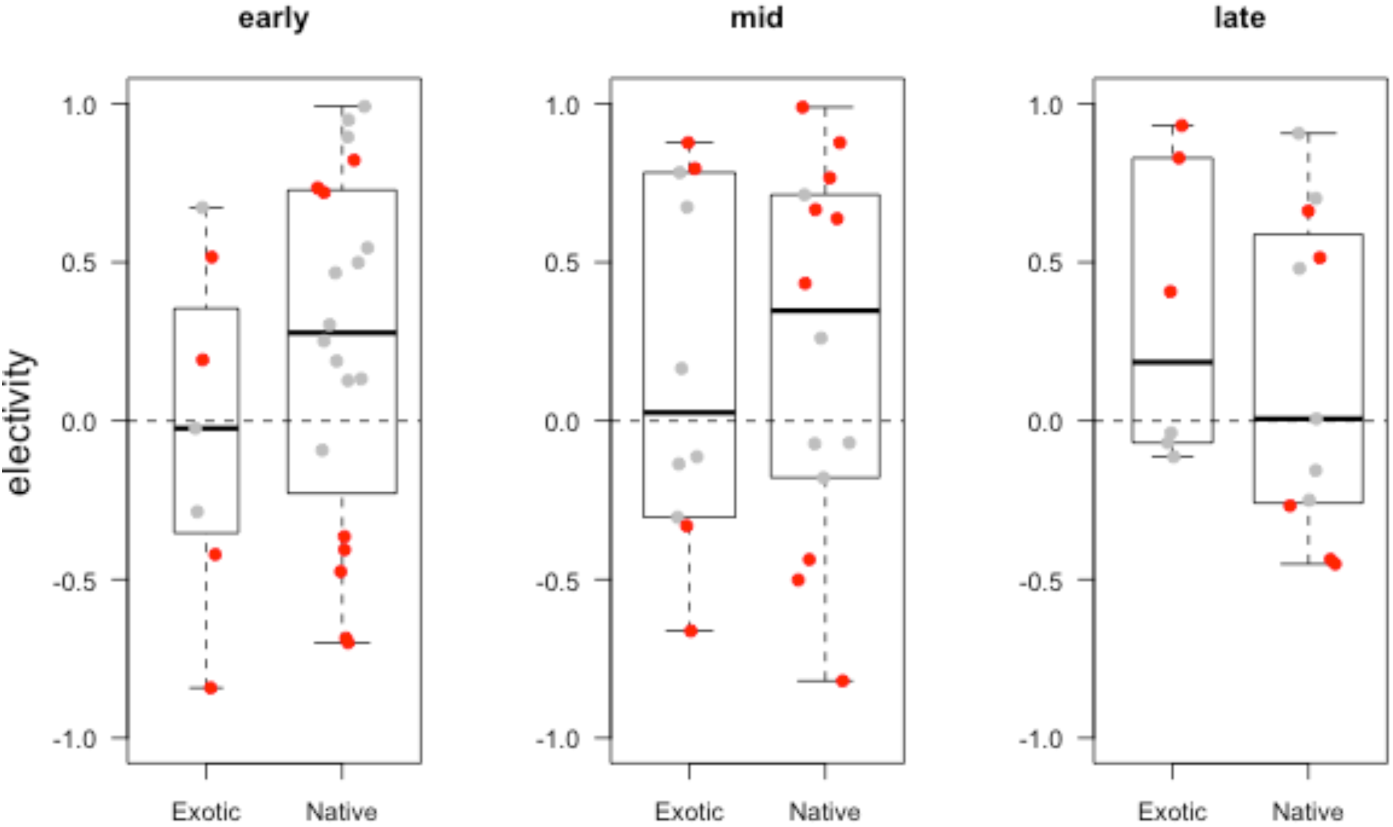
Boxplots of the *electivity* indices per plant species separated by plant origin (exotic, native) and season (early, mid, late). Each plant value is plotted in the background in grey when preference is not significant and in red when significant (significance based on Chi square Pearson residuals test). Positive values indicate plants that are preferred and negative values those that are avoided.

Second, we shift the focus of our analysis to analyze pollinator species-specific preferences. We excluded pollinator species with less than 20 visits recorded per season in order to prevent confounding rarity with specialization (Blüthgen 2010). We end up with 16 pollinator species, some of them present in several seasons, making 22 pollinator-season combinations. Again, each of the 22 pollinator-season combination was evaluated using Chi square tests and *electivity* values were compared between exotic and native plants for each pollinator species using mixed models with season as random effect. All pollinators showed significant preferences for certain plant species (chi square p-values < 0.05). When analyzed individually most pollinators do not show a consistent preference for exotic or native plants, with the exception of three species (Fig. 3; all species p-values > 0.1, except *Bombus melanopygus* = 0.05, *Dialictus incompletum* = 0.07 and *Halictus ligatus* = 0.03), all of them preferring exotics over natives. However, some pollinators of the 16 analyzed do prefer only one or more native plants (4 species: *Evylaeus sp, Osmia nemoris, Ceratina arizonensis, Calliopsis fracta*), others prefer only one or more exotic plants (7 species: *Synhalonia actuosa, Synhalonia frater, Bombus vosnesenskii, Halictus ligatus, Halictus tripartitus, Megachile apicalis, Bombus californicus*) and some prefer a mix of exotic and natives (4 species: *Bombus melanopygus, Ashmeadiella aridula, Dialictus incompletum, Dialictus tegulariforme*). The differential preferences regarding the exotic status create the basis for expecting winners and losers after an invasion process (see Fig. 1).

**Figure 3.**
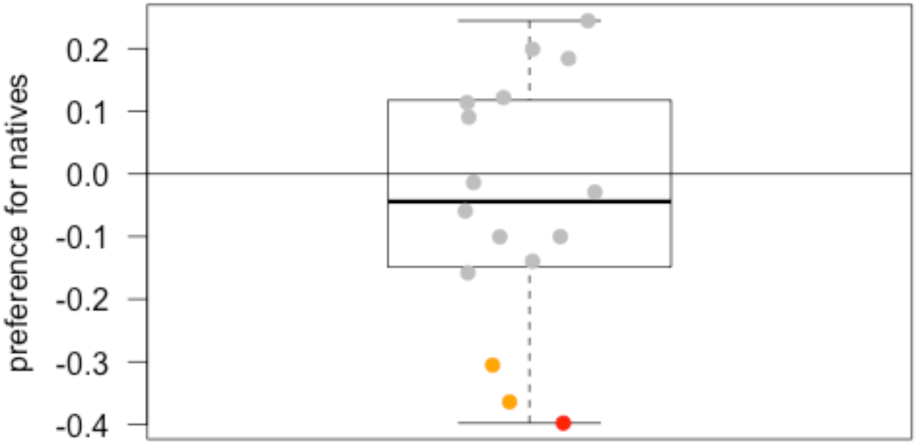
Boxplot of the effect sizes (i.e. model estimates) indicating the difference between *electivity* to exotics and to natives for each pollinator species. Positive values indicate overall preferences for natives, and negative values to exotics. Data points for each pollinator species are indicated in grey when not significant, in orange when p < 0.1 and in red when p < 0.05.

In conclusion, although the overall pattern is no preference for exotic plants, some particular exotic (and native) plants are overall preferred. Similarly, most pollinators do not have overall preferences for exotics, but a few species do favor them. Those are social species, usually common and sometimes even considered species typical of disturbed areas (e.g. *Halictus ligatus, Dialictus incompletum*). Interestingly, even within the species with no overall preference for exotics, we identify pollinators that prefer particular exotic plants. These pollinators are more likely to be positively affected by the invasion process, the others negatively affected, as their preferred resources will potentially diminish through displacement by invasive plant species.

## Relevance for conservation

We highlighted that pollinator species vary in response to plant invasions, including pollinators use, preference, and in some cases population dynamic consequences. Assessing the winners and losers in front of the rapid rate of invasive species introductions is crucial for understanding the responses of species groups performing important ecosystem functions, like pollination (Ollerton et al. 2011, Klein et al. 2007). There has been a recent awareness of pollinator declines globally (Potts et al. 2010, Gonzalez-Varo et al. 2013). However, biological invasions, especially by plant species, have received little attention as a threat (but see Stout and Morales 2008, and Morales et al. 2013 for effects of animal invasions). We already know that not all pollinators are equally affected by global change, a few are winners and many are losers (Bartomeus et al. 2013). Interestingly, among the winner pollinators we found species that are able to use flowering crops and tolerate new human-modified habitats (Bartomeus and Winfree 2013). Gaining information about which species are able also to exploit new exotic plants will be a way forward to understand which species will be flexible enough to survive in novel ecosystems, often dominated by exotic plants. In a changing world, species able to adapt their foraging strategies to use new resources may be the better suited to survive. For example, bumble bees that use the widespread plant invader *I. glandulifera* in the EU are thriving, while endangered bumble bee species do not use it (Kleijn and Raemaker 2008).

If we are going to manage emerging novel ecosystems, we need to incorporate pollinator specific responses to different global change drivers, including plant invasions. Some bumble bees and other trophic generalist bees can benefit from exotic plant invasions, as shown by the fact that those can use and even prefer to forage on new exotic plants. This behavioral flexibility may be the key to persisting in a changing world, and maintaining an important ecosystem function. More research is needed on the degree that plant invasions negatively affect those species in comparison with other disturbances that are occurring simultaneously. We need to implement better population monitoring programs at the community level (so indirect responses can be accounted for), but overall, understanding better which role play the pollinator behavior flexibility and cognitive capabilities in the process of adapting to novel environments is a promising line of research.

## Acknowledgments

We thank the book editors and an anonymous reviewer for their comments on the manuscript. IB acknowledge funding from the project SURVIVE_HIREC (CGL2013-47448-P)

